# Effects of Oxytocin Receptor Blockade on Dyadic Social Behavior in Monogamous and Non-Monogamous *Eulemur*

**DOI:** 10.1101/2022.09.14.507945

**Authors:** Nicholas M. Grebe, Alizeh Sheikh, Laury Ohannessian, Christine M. Drea

## Abstract

A prominent body of research spanning disciplines has been focused on the potential underlying role for oxytocin in the social signatures of monogamous mating bonds. Behavioral differences between monogamous and non-monogamous vole species, putatively mediated by oxytocinergic function, constitute a key source of support for this mechanism, but it is unclear to what extent this hormone–behavior linkage extends to the primate order. In a preregistered experiment, we test if oxytocin receptor blockade affects affiliative behavior in mixed-sex pairs of *Eulemur*, a genus of strepsirrhine primate containing both monogamous and non-monogamous species. Inconsistent with past studies in monogamous voles or monkeys, we do not find confirmatory evidence in *Eulemur* that monogamous pairs affiliate more than non-monogamous pairs, nor that oxytocin receptor blockade of one pair member selectively corresponds to reduced affiliative or scent-marking behavior in monogamous species. We do, however, find exploratory evidence of a pattern not previously investigated: simultaneously blocking oxytocin receptors in both members of a monogamous pair predicts lower rates of affiliative behavior relative to controls. Our study demonstrates the value of non-traditional animal models in challenging generalizations based on model organisms, and of methodological reform in providing a potential path forward for behavioral oxytocin research.

## Introduction

Animal groups vary greatly in their modal patterns of social organization, with one prominent source of variation expressed via strategies for mating and rearing offspring. Rather than engaging in transient mating relationships that dissolve after a reproductive cycle or mating season, mixed-sex pairs might instead live within a common home range and form long-term social relationships, an arrangement often referred to as social monogamy (Fernandez-Duque et al., 2020). Such an arrangement is often, though not always, accompanied by some degree of sexual exclusivity (“genetic monogamy”) or psychoaffective behavioral signatures (“pair-bonding”). Despite ongoing debate regarding how best to operationalize and separate the overlapping, yet distinct, dimensions that fall under this conceptual umbrella of ‘monogamy’ (e.g., Tecot et al., 2016; Bales et al., 2021), it is widely agreed that such social arrangements among sexual partners are relatively rare in mammals (∼5% of species), and somewhat more common, yet still atypical, in primates (29% of species; Lukas & Clutton-Brock, 2013). Here, we investigate the neuroendocrine underpinnings of such relationships in an experimental comparison of monogamous and non-monogamous strepsirrhine primates within the *Eulemur* genus.

In foundational work establishing new rodent models, researchers identified a potentially key role for the neuropeptides oxytocin and vasopressin in explaining the emergence of social monogamy and/or pair-bonding. Species within the *Microtus* genus of voles differ markedly in their mating systems: *M. ochrogaster* (prairie voles) form monogamous bonds characterized by strong partner preferences, whereas polygynandrous *M. montanus* and *pennsylvanicus* (montane and meadow voles, respectively) do not form such bonds and generally mate promiscuously (Insel & Shapiro, 1992). From this observation, researchers carried out various studies on these vole species with the goal of identifying a biological foundation of monogamy. For instance, they established that oxytocin receptor density differs substantially between monogamous and promiscuous vole species (Insel & Shapiro, 1992), that central administration of oxytocin mediates partner preference formation in prairie voles (Williams et al., 1994), and that oxytocin receptor blockade prevents partner preference formation in this same species (Cho et al., 1999; but see Berendzen et al., 2022). Neuroethological and neuroanatomical studies of the prairie vole have fostered a dominant paradigm for investigating the biological mechanisms underlying pair-bond formation and maintenance (reviewed in Walum & Young, 2018).

The promise and excitement of this work propelled an avalanche of studies testing the effects and/or correlates of oxytocin in human social bonds (reviewed in Gangestad & Grebe, 2016; Mierop et al., 2020). Despite a clear desire among researchers to translate insights from rodent models to human social behavior (see e.g. Young & Wang, 2004), patterns emerging from literature searches show a relative dearth of studies and experiments on nonhuman primates that would act as crucial intermediates in the translational gap between rodents and humans (Freeman & Bales, 2018; Grebe et al., 2021). The much smaller evidence base available on ‘monogamous’ primate species (including pair-housed monogamous or polyandrous callitrichids: Goldizen, 1988), in which researchers have examined if oxytocin-or vasopressin-mediated neurobiological mechanisms might underlie the emergence of monogamous social systems in primates, has yielded some positive support, albeit with numerous caveats and open questions.

For instance, in mixed-sex pairs of black-pencilled marmosets (*Callithrix penicillata*), oral administration of an oxytocin receptor antagonist (L-368,899; hereafter “OTA”) to one member of a mixed-sex pair decreased rates of proximity seeking and food sharing, had no significant effects on mating or sexual solicitation behavior, and had mixed effects on preferences for partners over strangers (Smith et al., 2010). In two other studies, conducted on mixed-sex pairs of common marmosets (*Callithrix jacchus*), researchers similarly found that OTA administration in one pair member led to decreases in proximity-seeking behavior during a stressor (Cavanaugh et al., 2016) or following an experimental separation (Cavanuagh et al., 2018). In cotton top tamarins (*Saguinus oedipus*), circulating oxytocin concentrations were reported to be related to social behavior in a sex-specific fashion, with receiving contact predicting concentrations only in females, and sexual activity predicting concentrations only in males (although it is unclear if sex-specific interactions reached statistical significance; see Snowdon et al., 2010). Finally, in another study on common marmosets, researchers found that administering oxytocin to individuals predicted how often they *received* grooming and proximity-seeking behavior from their partners, but not how often they *initiated* the same behavior; moreover, administering an OTA to a pair member did not lead to any comparable opposing behavioral changes (Cavanaugh et al., 2015).

In sum, although multiple findings are generally consistent with a role for oxytocin in nonhuman primate mating and pair-bonding, they are based on a variety of operationalizations, manipulations, and subgroup analyses, and researchers often report heterogeneous effects (see also Mustoe et al., 2018). It is thus difficult to draw clear conclusions about the hormone’s core functions in this domain. Indeed, a series of high-profile failures to replicate published findings on the hormone’s putative psychobehavioral effects in humans (e.g., trust: Nave et al., 2015; empathy/memory: Tabak et al., 2019; social interaction in children with autism: Sikich et al., 2021) has spurred recent scrutiny of oxytocin research more generally (see Mierop et al., 2020). Additionally, heterogeneity both within and between animal models regarding neuropeptide-mediated behavior (e.g., Madrid et al., 2020; Berendzen et al., 2022) and neurocircuitry (Phelps & Young, 2003; Grebe et al., 2021) underscores the need for robust, comparative approaches to both establish generalizable principles and identify disparate features of oxytocin’s social functions.

In the present study, we address several outstanding issues in oxytocin research. At a conceptual level, we combine the translational value of a primate model with the power of a ‘natural experiment’ of interspecific mating system variation. Namely, we investigate the effects of oxytocin system manipulation in regulating dyadic bonds of closely related monogamous and non-monogamous lemur species. We focus on the *Eulemur* genus of Malagasy primates, which is the sole primate analogue to *Microtus* voles in terms of mating system diversity within a single genus (see also Grebe et al., 2021). Behavioral and genetic evidence supports a structure of social monogamy and, potentially, pair-bonding in *E. mongoz* and *E. rubriventer* (mongoose and red-bellied lemurs, respectively). Male-female pairs in these two species typically live year-round in small family groups and defend a shared territory, engage in behavioral repertoires of mutual scent marking and vocalizations, and jointly care for young across several seasons; in the better-studied *E. rubriventer*, researchers have additionally found evidence for genetic monogamy (Curtis & Zaramody, 1999; Jacobs et al., 2018; Singletary & Tecot, 2019). All other species of *Eulemur*— as well as the ring-tailed lemur (*Lemur catta*), a close relative to this genus that we also included in our study—are non-monogamous, live in larger social groups, and show varying degrees of promiscuous mating (Kappeler & Fichtel, 2016). These features situate *Eulemur* as a powerful model for describing oxytocinergic functions unique to monogamous primates. At a methodological level, we perform a preregistered experiment (see https://osf.io/47mx3/) to cleanly distinguish between confirmatory effects of oxytocin system manipulation and exploratory, follow-up analyses that might help identify additional effects of oxytocin system manipulation.

## Methods

### Preregistered Confirmatory and Exploratory Predictions

Our preregistered predictions, experimental protocols, target sample size, exclusion criteria, and planned analyses are available on the Open Science Framework at https://osf.io/47mx3/. Any deviations from preregistered procedures or confirmatory analysis plans are explicitly noted in the sections below. We tested three confirmatory predictions that directly stem from previous research on monogamous rodents and primates and the hypothesized effects of oxytocin within these species (reviewed in e.g. Insel & Young, 2001; French et al., 2018). First, we examined rates of behavior in control conditions, expecting monogamous pairs to show higher average rates of affiliative behavior (allogrooming and time spent in physical contact) than non-monogamous pairs (following comparisons of vole species with different mating systems; e.g. Shapiro & Dewsbury, 1990; Salo et al., 1993). Second, we predicted oxytocin receptor blockade to affect the same affiliative behavior in individuals from monogamous, but not non-monogamous, pairs—reviews of the vole literature suggest that oxytocinergic effects on mating bonds are confined to monogamous pairs (e.g. Insel & Young, 2001, and studies of pair-living callitrichids demonstrate comparable effects (e.g. Smith et al., 2010). Thus, for this prediction we tested for an interaction term between mating system and OTA status in predicting our target social behavior. Third, we predicted that on days when monogamous individuals (but not non-monogamous individuals) are exposed to the scent of a potential reproductive competitor during a bioassay, oxytocin receptor blockade should diminish their rates of both scent marking and affiliative behavior toward partners relative to controls. This prediction is based on findings of oxytocin increases in humans (Grebe et al., 2017) and increases in affiliative behavior in pair-living callitrichids (Washabaugh & Snowdon, 1998) following the introduction of pair-bond threats. Here, we also examined an interaction between mating system and oxytocin receptor blockade, with this interaction specific to ‘opposite-sex bioassay’ days.

Next, we examined additional potential effects of OTA administration that do not rise to the level of confirmatory predictions due to absent and/or ambiguous patterns in previous research. Specifically, for each mating system, we compared rates of behavior across four experimental conditions (control, OTA administration to the female partner, OTA administration to the male partner, and OTA administration to both partners concurrently), as an elaboration on our confirmatory analyses that use a simple binary classification of whether or not a focal individual received the OTA on a given experimental day (i.e., ‘actor effects’ of OTA administration). Exploratory analyses allow us to examine a) potential ‘partner effects’—i.e., if modulating an individual’s oxytocinergic activity predicts their partner’s behavior—which received inconsistent support in a previous study of marmosets (Cavanaugh et al., 2015); and, relatedly, b) joint effects of OTA administration—i.e., if simultaneous OTA administration in both partners has behavioral effects distinct from ‘single-dose’ conditions in each sex. To the best of our knowledge, despite researchers using numerous varieties of oxytocin system manipulation in primates, no study has yet tested for potential joint effects of OTA administration. Previous findings led us to expect actor effects, rather than partner effects and/or joint effects, to explain behavioral variation, but we designed our study to explore additional possibilities via comparisons across all experimental conditions (as described under subheading b-iii on p. 2 of our preregistration document).

### Animals and Housing

Our study included 26 adult lemurs (22 *Eulemur spp*. and 4 *L. catta*; mean ± SD age: 15.6 ± 8.8 years), representing seven species, all tested as 13 well-established (i.e., successfully cohabitating > 6 months), mixed-sex pairs (Table 1). Six study pairs were either *E. mongoz* or *E. rubriventer* and thus designated as ‘monogamous’; the remaining seven pairs were designated as ‘non-monogamous’ (Table 1). The pairs were housed in one of three facilities: the Duke Lemur Center or DLC (n = 9) in Durham, North Carolina, USA; BioParc Valencia (n = 2) in Valencia, Spain; and the Lyon Zoo (n = 2), in Lyon, France (Table 1). Our study was originally planned for DLC animals only; however, owing to natural animal mortality during the course of the study, we ultimately expanded the number of facilities.

**Table 1.** Number of mixed-sex lemur pairs that participated in the present study, by mating system, species, and research facility.

| Facility | Monogamous lemur pairs |  | Non-monogamous lemur pairs |  |  |  |  |
| --- | --- | --- | --- | --- | --- | --- | --- |
|  | *E. mongoz*<br>(mongoose lemur) | *E. rubriventer*<br>(red-bellied lemur) | *E. collaris*<br>(collared lemur) | *E. coronatus*<br>(crowned lemur) | *E. flavifrons*<br>(blue-eyed black lemur) | *E. rufus*<br>(red-fronted lemur) | *L. catta*<br>(ring-tailed lemur) |
| Duke Lemur Center | 2 | — | 1 | 2 | 1 | 1 | 2 |
| BioParc Valencia | 1 | 1 | — | — | — | — | — |
| Lyon Zoo | — | 2 | — | — | — | — | — |

Animal housing, husbandry, and dietary conditions across institutions followed the AZA *Eulemur spp*. Care manual (AZA Prosimian Taxon Advisory Group, 2013); latitude was similarly comparable between institutions. In the Northern Hemisphere, the breeding season for these species ranges primarily from November – January (AZA Prosimian Taxon Advisory Group, 2013); we completed experimental procedures outside these months. Of the 13 pairs, 12 lived without additional enclosure-mates; one pair lived with a juvenile son, who was temporarily separated from his parents during experimental procedures and immediately reunited afterwards. Due to constraints on eligible lemur pairs, we stopped data collection after achieving our minimum planned sample size (at least six pairs from each mating system group) and did not perform any interim data-peeking. All procedures were approved by the Institutional Animal Care & Use Committee at Duke University, and veterinary staff at BioParc Valencia and the Lyon Zoo.

### Study design

Lemur pairs participated as a unit in a repeated measures experiment. Our primary manipulation, OTA administration (see below), consisted of the following four conditions, conducted in a randomized order per pair: (1) “Control” condition, in which pairs were observed absent any pharmacological manipulation; (2) “Female OTA” condition, in which only the female member received the antagonist; (3) “Male OTA” condition, in which only the male member received the antagonist); and (4) “Both OTA” condition, in which both members of the pair received an OTA. Each experimental condition lasted for a block period of three consecutive days; therefore, each animal was scheduled to receive the OTA for a total of six days.

Within each block, during two of the three daily observations that followed antagonist administration, we additionally probed the animals’ behavior by presenting an olfactory bioassay that proxied either the threat of a potential intruder or mating competitor. Specifically, we exposed pairs to the scent of a female stranger (“Female Bioassay”) or to the scent of a male stranger (“Male Bioassay”). On the remaining day, we presented no scent (“No Bioassay”). Pairs participated in one bioassay condition per day, with the order also randomized. Following experimental blocks, animals underwent a minimum washout period of five days between drug conditions (modified from the planned one-week washout in the preregistration document to accommodate staffing schedules; evidence suggests that a five-day period provides sufficient time for the OTA to be cleared; Boccia et al., 2007).

### Oxytocin Receptor Antagonist (OTA)

On experimental days, we temporarily blocked lemurs’ central and peripheral oxytocin receptors via oral administration of L-368,899, a pharmaceutical-grade OTA manufactured by MedChem Express (Monmouth Junction, NJ). L-368,899 is a non-peptide antagonist with a highly specific affinity for oxytocin receptors; it is reported to reliably cross the blood-brain barrier and persist in both the cerebral spinal fluid and periphery for several hours after administration (Boccia et al., 2007). During the morning hours (0700–1100 h), after morning feeding, we administered L-368,899 at a dose of 20 mg/kg—this dose followed previous studies, including Smith et al., (2010), Cavanaugh et al. (2016), and Cavanaugh et al. (2018). Given available evidence on the oral bioavailability of L-368,899 (16 – 41%; Thompson et al., 1997), we anticipated this dosage would meet or exceed that reported to generate behavioral effects via intravenous administration in rhesus macaques (*Macaca mulatta*; Boccia et al., 2007). The lemurs received the OTA infused into a preferred, hand-fed food item.

Individuals not receiving OTA, whether as part of treatment or control conditions, were fed preferred food items at the same time, and in the same manner, as treated animals. The individuals accepted the OTA during all six days scheduled, except for two females that refused to consume the antagonist partway through a condition block and only provided partial data for that block (see *Exclusions* below).

### Observations and Bioassays

Beginning 90 min after administration (or 90 min after feeding during control blocks), four trained observers performed 60-min focal observations on both members of the pair, simultaneously. During the observation sessions, we recorded select solitary behavior, including bouts of self-grooming, feeding, and all occurrences of scent marking (of either the environmental substrates or the partner, i.e., allomarking). We also recorded all dyadic interaction, including allogrooming, huddling, and aggression, and obtained a running count of the total time per observation that a pair spent in contact. We provide an ethogram of our target behavior in the Supplementary Online Materials (SOM). All behavior was recorded using the Animal Observer application (Caillaud, 2017). Most observations (73%) were scored concurrently in-person; the remaining 27% were video-recorded to be scored later in instances when there was no observer present who was blind to experimental condition. Additionally, each observer’s first in-person session was also video-recorded as a quality-control measure. An additional rater scored a subset of these videos to confirm inter-observer reliability; agreement on behavioral rates exceeded 85%.

During bioassays (representing 2 of 3 days per experimental block), the pairs were exposed to two wooden dowels placed at the edge of their enclosures at the beginning of the observation session. One dowel was rubbed with a sterile cotton swab, whereas the other dowel was anointed with the scent of an unrelated and unknown male or female conspecific. These ‘donor scents’ had been collected on pre-cleaned cotton swabs from the anogenital or genital glands of *Eulemur* and ring-tailed lemurs, respectively, during the breeding seasons of 2010-2019 (following previously published procedures; e.g., Greene & Drea, 2014). They had been frozen immediately and stored at -80 °C, until being thawed and applied to the dowels for our study. The left-right ordering of scented vs. unscented dowels alternated between study days. In addition to all behavior described above, we recorded occurrences of lemurs sniffing, licking, and scent marking each dowel. In analyses, we summed counts of olfactory behavior to assess engagement with the scented dowel, adjusting for engagement with the unscented dowel.

### Exclusions

There were two instances in which an animal refused to take the OTA. Our preregistered exclusion criteria specified these observations would be excluded from analyses, but in these cases, a female completely refused the drug during the “Both OTA” condition; because the male had already consumed his dose, we decided to deviate from our plan and treat the sessions as additional “Male OTA” experimental days. These two females did not receive the OTA on any days following their refusal. Additionally, there were three sessions (including the two individuals who would later refuse the drug entirely) in which an individual took approximately half of their antagonist dose, but refused the rest. We did not preregister a plan for such a scenario, but given these individuals received at least a partial dosage, we decided to retain these observations under their originally assigned conditions (see our Supporting Online Material [SOM] for robustness analyses excluding these observations, which yielded very similar results to those presented below).

### Data Analysis

We conducted all analyses in R (version 4.1.2) using multilevel models of individuals nested within dyads (using the *glmmTMB* package; Brooks et al., 2017); from these models, we subsequently report figures, marginal means, and contrasts produced by the *emmeans* package (Lenth, 2022). All code and data necessary to reproduce our results are publicly available on our OSF repository (https://osf.io/q2cvf/). Our statistical models predicted rates of affiliative behavior (allogrooming, time spent in contact) or scent marking from mating system, experimental condition (in confirmatory analyses, whether or not a focal individual received the OTA on that day; in exploratory analyses, pairwise comparisons between Control, Female OTA, Male OTA, and Both OTA), and their interaction. Following our preregistration, all models included age and sex as covariates and estimated separate variance structures for males and females. We modeled count outcomes (allogrooming, scent marking) with a negative binomial family and continuous outcomes (time spent in proximity) with the Tweedie family (for the latter, we deviated from our preregistered analysis plans after encountering convergence issues with the default Gaussian family). In all cases, we verified model fit by inspecting the deviation, dispersion, and outliers of residuals using the *DHARMa* package (Hartig, 2022). We describe statistical significance in both our confirmatory and exploratory analyses, acknowledging that the latter cannot be interpreted in the same manner as the former. For comparison with confirmatory results, we also report and emphasize effect sizes in Cohen’s *d* in exploratory analyses.

## Results

### Confirmatory Analyses

Contrary to our first confirmatory prediction that, absent any antagonist administration, monogamous pairs would engage in higher rates of affiliative behavior than would non-monogamous pairs, we found no significant differences in behavior as a function of mating system. Comparisons revealed that model-estimated rates of affiliative behavior were actually modestly, though not significantly, higher in non-monogamous pairs: for time huddling, 35% versus 22% (*t*(149) = 1.79, *p* = 0.075, *d* = 0.29; Figure 1A); for hourly bouts of allogrooming, 1.66 versus 0.99 (*t*(150) = 1.14, *p* = 0.258, *d* = 0.19; Figure 1B).

**Figure 1.**
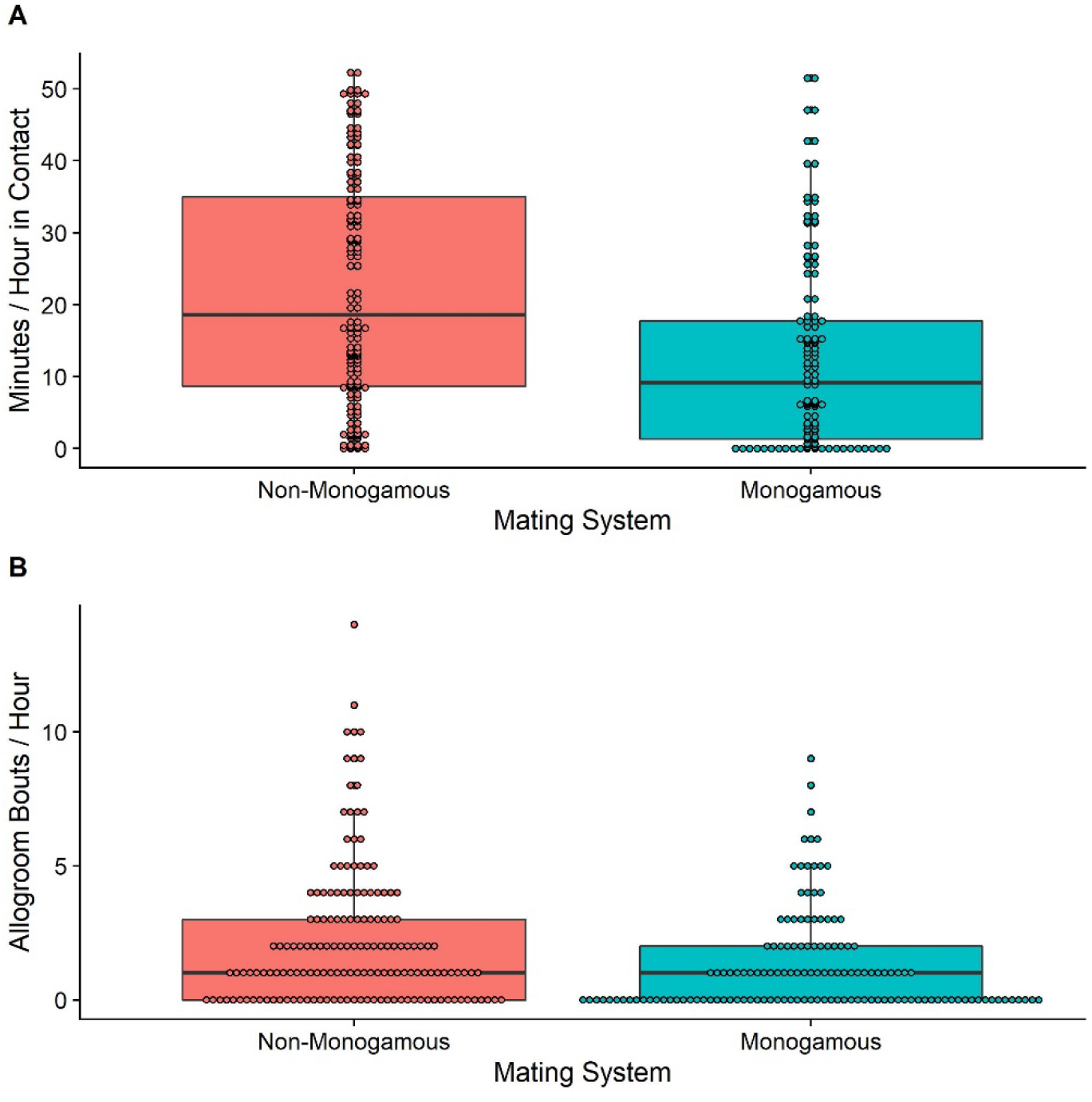
Box and dot plots showing rates of (A) close contact and (B) allogrooming during control observations of mixed-sex pairs of non-monogamous and monogamous lemurs.

In our second set of confirmatory analyses, we predicted an interaction between mating system and actor experimental condition. This interaction was non-significant for allogrooming (*z* = -1.62, *p* = 0.104), and the contrast between drug and control conditions did not significantly differ from zero in dyads of either mating system (*t*(287) = -1.17, *p* = 0.242 and *t*(287) = 1.16, *p* = 0.246 for non-monogamous and monogamous pairs, respectively). This interaction fell just short of significance for time spent huddling (*z* = -1.84, *p* = 0.067). Decomposing this latter interaction into simple effects showed that blocking oxytocin receptors did not change contact time in non-monogamous pairs (*t*(287) = -0.10, *p* = 0.920, *d* = -0.01), but did lead to a moderate decrease in contact time in monogamous pairs (*t*(287) = 2.27, *p* = 0.024, *d* = 0.27); nevertheless, these simple effects did not significantly differ from one another.

In our third set of confirmatory analyses, we also predicted an interaction between mating system and actor experimental condition: here, for individuals in monogamous pairs only, we expected changes in affiliative and scent-marking behavior during the presentation of scents from potential competitors (i.e. during opposite-sex bioassay conditions). For time spent huddling, the target interaction was not significant (*z* = -1.07, *p* = 0.284); there was no evidence that blocking oxytocin receptors specifically affected huddling in monogamous pairs on bioassay days. The target interaction for allogrooming fell just short of significance (*z* = -1.86, *p* = 0.063); unexpectedly, decomposing this interaction into simple effects revealed that blocking oxytocin receptors led to marginally more allogrooming in non-monogamous pairs (*t*(84) = -1.77, *p* = 0.080, *d* = -0.39) and non-significantly less allogrooming in monogamous pairs (*t*(84) = 1.12, *p* = 0.264, *d* = 0.24). Lastly, there was no evidence of an interaction between antagonist administration and mating system on interest toward strangers’ scents during bioassay trials (*z* = -0.98, *p* = 0.328).

### Exploratory Analyses

We next investigated if there were potential a) partner effects and/or b) pairwise differences in target behavior between any of the four experimental conditions (Control, Female OTA, Male OTA, Both OTA). For affiliative behavior (huddling and allogrooming), we did not observe any significant interactions between partner experimental condition and mating system (all *p*s > 0.05); i.e., a monogamous individual receiving the OTA did not predict their partner’s affiliative behavior, when actor effects of the OTA were held constant. By contrast, we observed two similar, significant interactions between overall experimental condition and mating system, suggesting a synergistic OTA effect: affiliative behavior occurred at the lowest rates in the “Both OTA” condition of monogamous pairs, but not of non-monogamous pairs. Monogamous pairs huddled on average 23% of the time in control conditions, but only 11% of the time when both members were administered the OTA (Figure 2A). This contrast corresponded to a moderate effect size (*d* = 0.38). Rates of affiliative behavior in the “Female OTA” and “Male OTA” conditions were intermediate (22 % and 17%) and each significantly higher than in the “Both OTA” condition (*d* = -0.40 and -0.23; Figure 2B).

**Figure 2.**
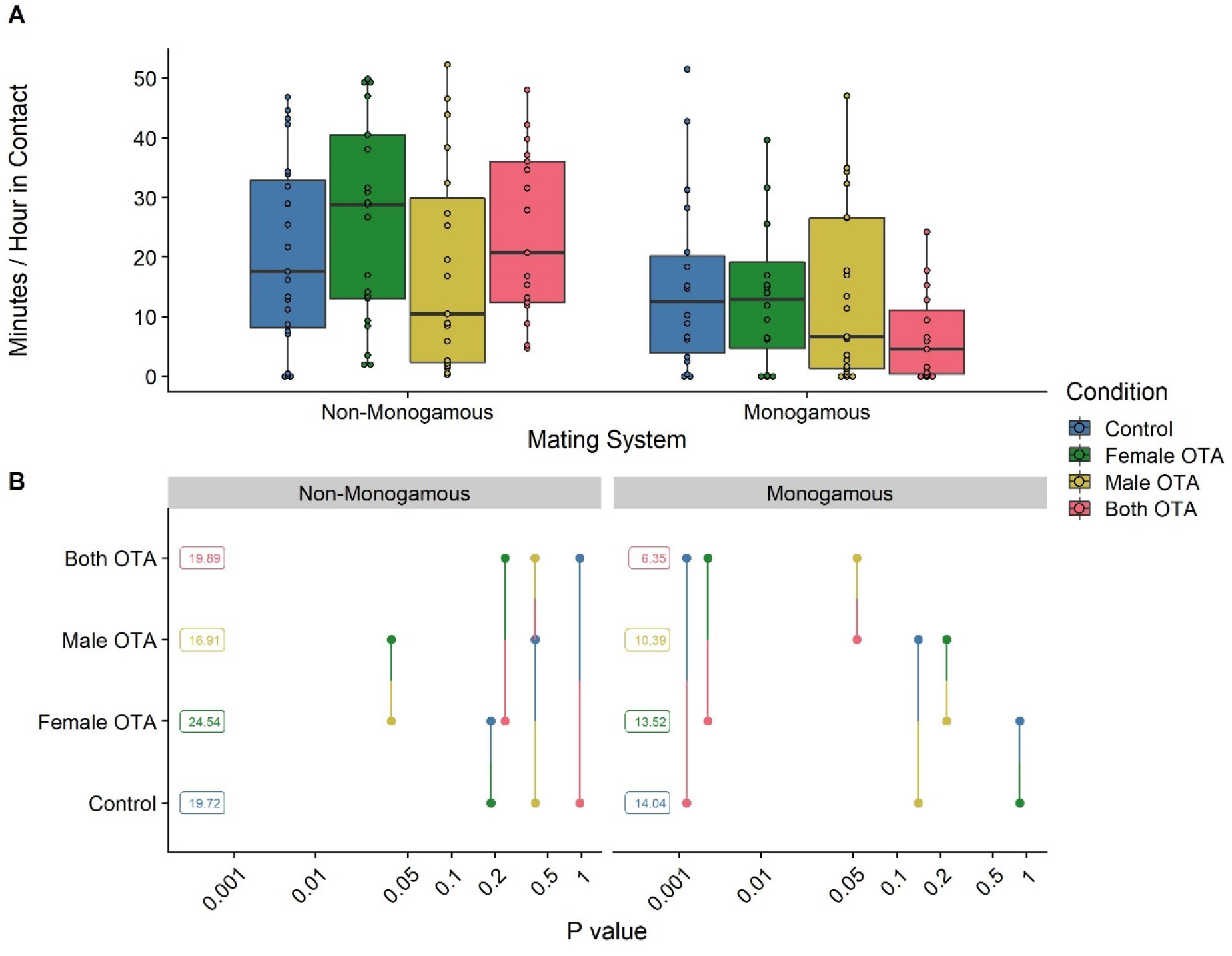
Lemur social contact in relation to oxytocin receptor treatment condition and mating system: (A) Box and dot plots comparing time spent in contact across the four, color-coded experimental conditions, with separate panels for each mating system; (B) pairwise p-value plots for comparisons within mating system. In pairwise *p*-value plots, factor levels are plotted on the vertical scale, and *p-*values are plotted on the horizontal scale. Each *p-*value is plotted twice at vertical positions, corresponding to the levels being compared and connected by a line segment (see Lenth, 2022).

For allogrooming, rates were once again lowest within monogamous pairs in which both partners had their oxytocin receptors blocked (0.46 bouts/hour; Figure 3A). Rates in the Both OTA condition were significantly lower than rates in the Control (0.99 bouts/hour; contrast *d* = 0.25) and Female OTA (1.50 bouts/hour; contrast *d* = 0.40) conditions, and were marginally less than rates in the Male OTA condition (0.91 bouts/hour; contrast *d* = 0.23) (Figure 3B).

**Figure 3.**
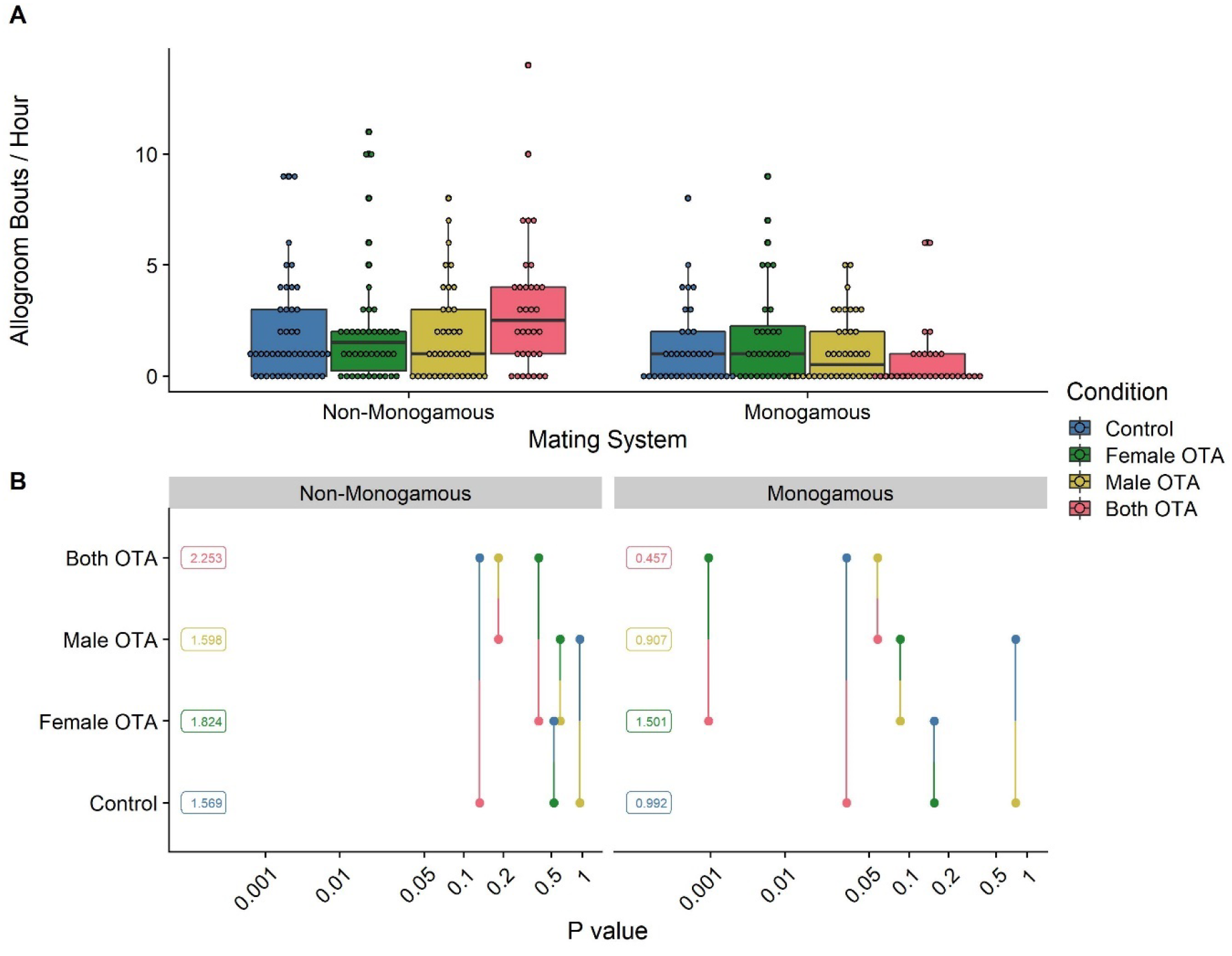
Lemur social grooming in relation to oxytocin receptor treatment condition and mating system: (A) Box and dot plots comparing bouts of allogrooming per hour across the four, color-coded experimental conditions, with separate panels for each mating system; (B) pairwise *p*-value plots for comparisons within mating system. In pairwise *p*-value plots, factor levels are plotted on the vertical scale, and *p-*values are plotted on the horizontal scale. Each *p-*value is plotted twice at vertical positions, corresponding to the levels being compared and connected by a line segment (see Lenth, 2022).

We also observed a significant interaction between experimental condition and mating system for interest in a stranger’s scent during the bioassay trials, although the form of this interaction differed from that observed for affiliative behavior. No contrasts between conditions differed substantially for non-monogamous pairs (all *p*s > 0.21, *d* < 0.22), whereas for monogamous pairs, interest in conspecific scents in the Male OTA condition was significantly lower than in either the Both OTA or Female OTA conditions (*d* = 0.46 and 0.65 for each contrast) and was non-significantly lower than in the Control condition (*d* = 0.34). We confirmed that this pattern was attributable to behavioral differences in males, as estimated effect sizes for the above contrasts were larger and of similar significance when re-running this statistical model in males only (Figure 4).

**Figure 4.**
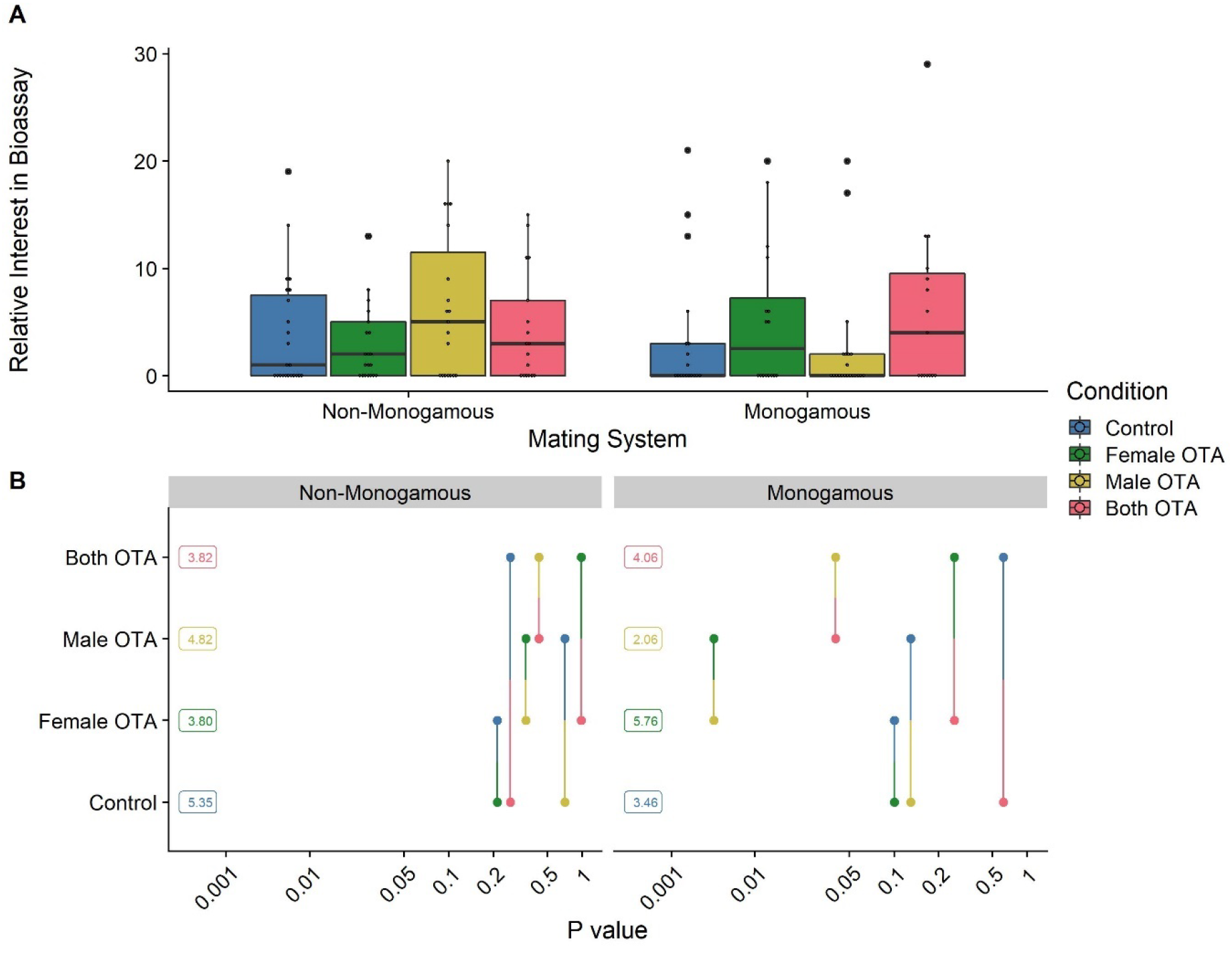
Lemur interest in conspecific scent in relation to oxytocin receptor treatment condition and mating system: (A) Box and dot plots comparing relative interest in the bioassay (interest in the scented dowel minus interest in the unscented dowel) across the four experimental conditions, with separate panels for each mating system; (B) pairwise *p*-value plots for comparisons within mating system. In pairwise *p*-value plots, factor levels are plotted on the vertical scale, and *p-*values are plotted on the horizontal scale. Each *p-*value is plotted twice at vertical positions, corresponding to the levels being compared and connected by a line segment (see Lenth, 2022).

## Discussion

The present study, conducted within the only primate genus known to contain both monogamous and non-monogamous species, provides new information about the effects and potential functions of oxytocin in mediating mating bonds and, more broadly, speaks to a number of open questions within contemporary oxytocin research and evolutionary primatology. Notably, contrary to predictions based on monogamous and non-monogamous voles (*Microtus* spp: Salo et al., 1993) socially monogamous *Eulemur* pairs did not show greater rates of affiliative behavior at baseline than did their non-monogamous counterparts. Additionally, expectations of an effect of OTA administration on affiliative behavior within monogamous *Eulemur* pairs, as occurs in callitrichids (Smith et al., 2010; Cavanaugh et al., 2016, Cavanaugh et al., 2018), had mixed support. Based on our restricted confirmatory analyses, we did not observe consistent evidence that administering an OTA to an individual within a monogamous pair led to less time in contact or grooming within the pair, nor did treatment lead to less overt interest in the scents of potential reproductive competitors. In exploratory analyses, however, we found that while disrupting a single individual’s oxytocin system produced no detectable changes in social behavior within monogamous pairs, simultaneously blocking both individuals’ oxytocin receptors did. Together, our findings suggest that variability across studies may relate to a) interspecific and/or higher-level taxonomic differences, b) pair-level heterogeneity in socially monogamous *Eulemur* mating bonds, and/or c) differences between effects of individual and joint OTA administration. This latter, novel finding should be addressed and tested in other study systems. Overall, the variability in our findings may also indicate d) an increasingly uncertain role of oxytocin in mating bonds (Grebe et al., 2021; Berendzen et al., 2022).

Our null results for baseline behavioral differences, and the additional absence of confirmatory evidence of behavioral changes following oxytocin receptor blockade, are subject to various potential interpretations. At one extreme, one might view the rates of affiliative social behavior displayed by monogamous and non-monogamous *Eulemur* pairs, with or without OTA treatment, as evidence that these species lack a psychological or behavioral pair-bond, as conventionally defined (see e.g., Bales et al., 2021). We recognize that more structured tests of the affective aspects of a pair-bond (e.g., separation distress, partner preference, and stress buffering; Bales et al., 2021) are lacking in *E. rubriventer* and *E. mongoz*, and that much less is known about the dynamics of long-term reproductive bonds in *Eulemur* than in other primate models (e.g., titi monkeys: Bales et al., 2017; marmosets and tamarins: French et al., 2018). We designed our experiment to detect effects of oxytocin system manipulation, rather than to provide precise characterizations of *Eulemur* bonding behavior and interspecific variation at baseline, but further investigation of the latter is clearly needed. Within the constraints of justifiable research on endangered species, further investigation of lemur social bonding, both in captivity and in the wild, will build upon initial investigations (e.g. Singletary & Tecot, 2019) and provide better understanding of the nature of monogamy in this understudied clade.

For various reasons, however, we do not favor an interpretation that monogamy in lemurs is fully distinct in form and function from that observed in more popular animal models of monogamous mating (e.g., *Microtus* spp: Salo et al., 1993; coppery titi monkeys (*Plecturocebus cupreus*): Bales et al., 2017; marmosets (*Callithrix* spp.): Ågmo et al., 2012, French et al., 2018; gibbons (*Hylobates* spp.): Palombit, 1996, cf. Reichard et al., 2012). If anything, perhaps a higher ‘baseline’ of sociality in the primate order, compared to rodents, reduces the scope of interspecific differences in affiliation attributable to mating system variation (for critical discussions of primate social ‘exceptionalism’, see e.g., Rowell, 1999; Silk & Kappeler, 2017); unfortunately, we lack the non-monogamous cohort among anthropoid models that would provide the basis for comparatively assessing in primates our null baseline differences in *Eulemur*. Moreover, the better-studied patterns of wild *E. rubriventer* bonding behavior, relative to *E. mongoz*, might mask substantial heterogeneity that exists even between monogamous members of the same genus. As previously noted, both *E. mongoz* and *E. rubriventer* show evidence of social monogamy in the wild, and use mutual territorial marking, allomarking, and huddling in a manner consistent with behavioral manifestations of pair-bonding (Curtis & Zaramody, 1999; Singletary & Tecot, 2019; Singletary & Tecot, 2020). Additional preliminary evidence in *E. rubriventer* also suggests the use of unique multimodal communication patterns to foster or advertise mating bonds (Singletary & Tecot, 2019), the latter of which has been suggested for other bonded lemur species (Greene & Drea, 2014), though it is unknown if similar mechanisms exist in *E. mongoz*. Thus, it remains to be determined if manifestations of “bonded-ness” differ between monogamous lemur species, and/or if characteristics of lemur bonding are distinct from the general patterns observed across monogamous anthropoid primates or rodents. We argue that broadening the taxonomic representation of study species might require some modification of the definitional criteria or attributes used to assess pair-bonding.

*Eulemur* neuroanatomy provides additional contextualization for our findings. In a recent study, our research group published the first maps of central oxytocin and vasopressin receptors in several species of monogamous and non-monogamous *Eulemur* and found little evidence, based on either receptor, of a “pair-bonding circuit” specific to monogamous individuals (Grebe et al., 2021). Instead, we observed substantial heterogeneity in receptor distributions between individuals of the same mating system. Individual animals whose brains were analyzed in Grebe et al. (2021) represent a distinct sample from the current study of living pairs, so our results cannot be directly correlated; however, mixed neuroanatomical results may provide one possible explanation for mixed behavioral results in the present study. If individuals of a socially monogamous species vary substantially in their distributions and densities of oxytocin receptors—perhaps by virtue of variation in their social histories—this variability should have consequences for the effects of oxytocin receptor blockade on social behavior—an assertion supported by invasive studies of monogamous prairie voles (e.g., Ross et al., 2009; Ophir et al., 2012). In our study, differences in affiliative behavior between the Both OTA and Control conditions—i.e., one plausible measure of the ‘total’ OTA effect—showed substantial between-pair variation in monogamous species, ranging from almost no difference to a severalfold increase. One possibility is that, in our study, effects of the experimental antagonist on affiliative behavior towards a mate might be limited to individuals that possess a distribution of receptors more reminiscent of previously characterized animal models of pair-bonding (i.e., dense binding in dopaminergic regions of “pair-bonding circuits”; Young & Wang, 2004).

In sum, we cannot rule out the possibility that null results are to some degree attributable to sources of variation beyond evolved differences in *Eulemur* mating systems: this variation includes not just heterogeneity in ‘monogamous’ bonding and neuroanatomy, but also the use of multiple study sites and long-term pair housing for non-monogamous species (cf. Grebe et al., 2021). We note that this latter point also applies to other primate groups, such as callitrichids, that are pair-housed in captivity but frequently mate polyandrously in the wild (Goldizen, 1988). Despite the potential noise added by these factors, the most ‘global’ disruption of oxytocinergic functioning in our study, in which both members of the dyad received a receptor antagonist, selectively corresponded to the lowest rates of affiliative behavior in *E. mongoz* and *rubriventer*, an effect in line with the general hypothesis that oxytocin differentially regulates dyadic behavior within monogamous mating bonds. We transparently acknowledge that this pattern arose from exploratory analyses, rather than our confirmatory predictions based on previously published effects of OTA administration; nevertheless, given the rare opportunity to perform a hormonal manipulation in endangered primates, we designed a study to explore if partner effects and/or additive effects of joint OTA administration might predict differences in behavior. Because evidence of this latter pattern has not been previously investigated nor reported in OTA research, we find it worthwhile to speculate on potential explanations.

One natural question is why, among monogamous pairs, rates of dyadic behavior in either the Female OTA or Male OTA condition were not distinguishable from controls, whereas rates in the Both OTA condition were distinguishable. This divergent pattern may arise because a) “single dose” conditions result in smaller changes in affiliative behavior that are difficult to detect without larger sample sizes and/or b) oxytocin disruption in either single member of a monogamous pair can be compensated for by their partners’ behavior. Because we do not observe a consistent “intermediate” disruption of affiliative behavior in the Female and Male OTA conditions, our observed results appear to be more consistent with partner compensation. Cavanaugh et al. (2015) report a partner effect of oxytocin administration on proximity seeking in marmoset pairs, in which males treated with oxytocin, relative to controls, received greater proximity seeking from their untreated female partners. Although Cavanaugh et al. (2015) do not find evidence that the same OTA we used in the current study had a comparable opposing effect on partner behavior, their suggestion that modification of the oxytocin system might affect partner responses to subtle social signals provides the basis for one potential interpretation. Monogamous *Eulemur* pairs, like other monogamous primates, may possess a reciprocal preference to initiate physical and social contact with their partners (Ågmo et al., 2012; Bales et al., 2021). If this behavior is, in part, mediated by oxytocin—perhaps via its activity in increasing the social reward value of a mate’s odor cues (Keverne & Curley, 2004)—disruption in one member of a well-established dyad might diminish that individual’s initiation of behavior, but an untreated partner might simply compensate for that decrement by increasing how often it initiates affiliation. Blocking oxytocin in both members, by contrast, might more consistently result in less affiliation because neither member is motivated to initiate huddling or grooming. Psychologists using intranasal administration paradigms have suggested that oxytocin may help with partner synchronization in social tasks when a partner is unresponsive (Gebauer et al., 2016); in a variation on this argument, perhaps normal oxytocin functioning within at least part of a monogamous mating bond is sufficient to coordinate affiliative bonding behavior. Relatedly, in a provocative new preprint, Berendzen and colleagues (2022) present evidence in prairie voles that oxytocin-receptor null mutants show a robust partner preference and repertoire of pair-bonding behavior comparable to wildtype individuals. Interestingly, all mutant individuals in their study were paired with wildtype partners. Similar to results from our current study, their finding raises the possibility that unmanipulated partners might somehow compensate for individuals that lack normal oxytocinergic functioning.

More broadly, the mixed results in our current study can be seen as joining a decidedly scattered pattern of results in the field of behavioral oxytocin research, including within foundational domains, such as trust, social cognition, and pair-bonding (e.g., Nave et al., 2015; Mierop et al., 2020; Berendzen et al., 2022). Collectively, results increasingly do not produce a clear consensus on reliable, generalizable behavioral effects and/or correlates of oxytocin, nor on predictable patterns of lineage-specific variation. One potential path forward may come from the adoption of open science methods and practices, such as preregistration/registered reports (Chambers & Tzavella, 2022), the sharing of open data, and multi-site collaboration.

Preregistration of analytic choices and key predicted outcomes can help with establishing which of the many overlapping, yet distinct, reported effects of oxytocin and OTA administration are robust, either within or between species. In particular, we encourage efforts to replicate in other monogamous species the “double dose” OTA pattern in our current study. Relatedly, preregistered, multi-site studies that aggregate information from datasets spanning diverse species and research sites have been central to assessing commonality and diversity in primate cognition (see ManyPrimates, 2022); comparable large-scale analyses of oxytocin–behavior relationships in primates may be a similarly fruitful avenue for future study.

In sum, our study produces some evidence consistent with a role for oxytocin in regulating *Eulemur* monogamy, but it also shows that several assumed characteristics of monogamy and oxytocin-mediated behavior are challenged by research on a rare, understudied group of primates. Non-traditional animal models thus prove their value for investigating both commonality and diversity of function within behavioral endocrinology. We expect that broadening the range of animal models studied will continue to reveal unanticipated findings that require modifications to existing theories of social behavior and neuroendocrinology (Rosenthal et al., 2017; Freeman & Bales, 2018; Grebe et al., 2021).

## Acknowledgments

We are especially grateful to staff at the Duke Lemur Center, BioParc Valencia, and the Lyon Zoo, as well as Annika Sharma, Amika Ekanem, and Cati Gerique, whose assistance with logistics and data collection made this study possible. This research was supported by the Duke Lemur Center’s Director’s Fund and National Science Foundation (NSF) Grant SBE-1808803 (to N.M.G. and C.M.D.). The collection of odorants for bioassays was funded by NSF IOS-0719003 and BCS-1749465 (to C.M.D.), and NSF BCS-1341150 and the Margot Marsh Biodiversity Fund (to C.M.D. and J. Petty).

This is DLC publication # ____.

