## Supplementary Online Materials for "Effects of Oxytocin Receptor Blockade on Dyadic Social Behavior in Monogamous and Non-Monogamous *Eulemur*"

**Appendix 1. Results excluding experimental sessions (n = 3) where a treated individual only took a partial dose of the oxytocin receptor antagonist (OTA). Full model results available from data and code posted publicly at [https://osf.io/q2cvf/](https://osf.io/q2cvf/).**

**Table S1. Omnibus statistics for confirmatory analyses (i.e., actor effects only; see Methods). Effects $p$ < 0.05 bolded.**

| | Parameter | | | | |
|---|---|---|---|---|---|
| | **Age** | **Mating System** | **OTA Administration** | **Sex** | **Mating × OTA Admin** |
| **Huddling** | $\chi^2(1) = 0.26$, $p = 0.605$ | $\chi^2(1) = 0.69$, $p = 0.406$ | $\chi^2(1) = 0.02$, $p = 0.903$ | $\chi^2(1) = 0.01$, $p = 0.973$ | $\chi^2(1) = 3.39$, $p = 0.065$ |
| **Allogroom** | $\chi^2(1) = 0.17$, $p = 0.682$ | $\chi^2(1) = 0.17$, $p = 0.680$ | $\chi^2(1) = 1.72$, $p = 0.190$ | $\chi^2(1) = 0.19$, $p = 0.660$ | $\chi^2(1) = 2.92$, $p = 0.088$ |
| **Scent Interest** | **$\chi^2(1) = 9.67$, $p = 0.002$** | $\chi^2(1) = 1.16$, $p = 0.282$ | $\chi^2(1) = 0.21$, $p = 0.647$ | $\chi^2(1) = 0.22$, $p = 0.642$ | $\chi^2(1) = 0.88$, $p = 0.347$ |

**Table S2. Estimated marginal means and standard errors for confirmatory analyses.**

| | Monogamous | | Non-Monogamous | |
|---|---|---|---|---|
| | **No OTA** | **OTA** | **No OTA** | **OTA** |

| | | | | |
|---|---|---|---|---|
| **Huddling** | 12.81 (2.79) | 8.94(2.03) | 19.94 (3.95) | 20.25 (4.13) |
| **Allogroom** | 1.11 (0.36) | 0.81 (0.27) | 1.72 (0.52) | 1.95 (0.58) |
| **Scent Interest** | 2.61 (0.82) | 4.53 (1.32) | 4.81 (1.27) | 3.73 (1.02) |

**Table S3. Omnibus statistics for exploratory analyses (i.e., partner effects and joint effects; see Methods).**

| | **Parameter (Joint Effects)** | | | | | **Parameter (Partner Effects)** | |
|---|---|---|---|---|---|---|---|
| | **Age** | **Mating System** | **Experimental Condition** | **Sex** | **Mating × Condition** | **Partner Condition** | **Mating × Partner Condition** |
| **Huddling** | $\chi^2(1)$ =0.26, $p$ = 0.610 | $\chi^2(1)$ = 1.09, $p$ = 0.300 | $\chi^2(3)$ =3.46, $p$ = 0.326 | $\chi^2(1)$ =0.00, $p$ = 0.976 | $\chi^2(3)$ = 7.49, $p$ = 0.050 | $\chi^2(1)$ =0.00, $p$ = 0.969 | $\chi^2(1)$ =3.21, $p$ = 0.073 |
| **Allogroom** | $\chi^2(1)$ =0.23, $p$ = 0.632 | $\chi^2(1)$ = 0.98, $p$ = 0.322 | $\chi^2(3)$ =2.79, $p$ = 0.424 | $\chi^2(1)$ = 0.26, $p$ = 0.608 | $\chi^2(3)$ =10.91, $p$ = 0.012 | $\chi^2(1)$ =0.48, $p$ = 0.487 | $\chi^2(1)$ =2.39, $p$ = 0.122 |
| **Scent Interest** | $\chi^2(1)$ =8.76, $p$ = 0.003 | $\chi^2(1)$ =0.99, $p$ = 0.319 | $\chi^2(3)$ =2.32, $p$ = 0.509 | $\chi^2(1)$ =9.95, $p$ = 0.019 | $\chi^2(3)$ = 0.02, $p$ = 0.903 | $\chi^2(1)$ =1.36, $p$ = 0.243 | $\chi^2(1)$ =6.54, $p$ = 0.011 |

**Table S4. Estimated marginal means and standard errors for exploratory analyses.**

| | **Monogamous** | | | | **Non-Monogamous** | | | |
|---|---|---|---|---|---|---|---|---|
| | **Control** | **Female OTA** | **Male OTA** | **Both OTA** | **Control** | **Female OTA** | **Male OTA** | **Both OTA** |
| **Huddling** | 14.06 (3.32) | 13.55 (3.31) | 10.42 (2.50) | 6.36 (1.75) | 19.65 (4.16) | 23.95 (5.19) | 17.09 (3.81) | 20.03 (4.47) |
| **Allogroom** | 0.99 (0.35) | 1.50 (0.53) | 0.91 (0.32) | 0.46 (0.19) | 1.59 (0.51) | 1.84 (0.61) | 1.62 (0.53) | 2.30 (0.74) |

| | | | | | | | | |
|---|---|---|---|---|---|---|---|---|
| **Scent Interest** | 3.42 (1.11) | 5.68 (1.79) | 2.03 (0.72) | 4.02 (1.28) | 5.24 (1.49) | 3.48 (1.07) | 4.82 (1.37) | 3.70 (1.11) |

**Appendix 2. Ethogram for affiliative, aggressive, and scent-marking behavior recorded during focal observation of lemur pairs.**

| **Behavior** | **Description** |
|---|---|
| *Scent mark* | |
| scent-mark substrate | Rubs the scent glands of the wrists, head, or anogenital region on a surface |
| scent-mark partner | Rubs the scent glands of the head or anogenital region on an enclosure-mate |
| *Aggression* | |
| cuff | Actor uses hand and arm to aggressively swipe at recipient |
| lunge | Actor lurches toward opponent as if to attack |
| chase | Actor aggressively pursues partner |
| bite | Actor places mouth and teeth on partner, with aggressive force |
| fight bout | Multiple, fast, aggressive interactions between both individuals, with the actor being the initiator |
| *Affiliative* | |
| self-groom | Actor runs tongue or tooth-comb over own fur |
| allogroom | Actor runs tongue or tooth-comb over other animal's fur |
| huddle | Approaches other animal and rests in body contact |
